# Genetic and physical interaction of Drosophila Ino80 with Polycomb Responsive Element

**DOI:** 10.1101/793778

**Authors:** Mohsen Ghasemi, Jayant Maini, Shruti Jain, Vasanthi Dasari, Rakesh Mishra, Vani Brahmachari

## Abstract

The chromatin remodeling protein, dIno80 (*Drosophila* Ino80) regulates homeotic genes. We show that Ino80, along with Trx and ETP (Enhancer of Trithorax and Polycomb) proteins, interacts with two Polycomb/Trithorax Responsive Elements (PRE/TRE), *iab-7* and *bxd PRE* in flies and the larval imaginal discs. In S2 cells, dIno80 localizes to the endogenous *iab-7* and *bxd-PREs*. The localization of Ino80 and Pleiohomeotic (Pho) at the PRE is sensitive to the cellular abundance of each other; when levels of *Ino80* are limiting, there is increased Pho enrichment, and *Pho* knock-down leads to increased enrichment of Ino80. We demonstrate that over-expression of dIno80 rescues the pupal lethality in *pleiohomeotic* (*pho*) deficient flies, which suggests that dIno80 has a role in cellular memory. The apparent competition between Pho and Ino80 for binding at the PRE indicates that Ino80 may act as a potential recruiter of the regulatory complex in addition to being a chromatin remodeler.

**Author Summary:** The null mutants of Pho and dIno80 show lethality at different stages of development in the fly, implying that they may function independent of each other. The observation that Pho-lethality can be rescued by overexpression of dIno80 with significant penetrance and that Ino80 has its own DNA binding domain, led us to predict that Ino80 may have Pho-independent functions, perhaps through non-canonical complexes. In the current study, we show that dIno80 interacts with *bxd* and *iab-7* PRE in cooperation with Polycomb and Trithorax proteins and regulate the homeotic genes. The effect of knock-down or mutation of dIno80 results in altered phenotype in adult flies and rescue of Lac-Z expression in imaginal discs, in parallel with similar effect of Pho mutation or knock-down. We provide evidence of direct interaction of dIno80 with *iab7-* and *bxd*-PRE using chromatin immunoprecipitation. The dIno80 localization in and around the PRE sequence was enhanced in the absence of Pho, indicating competition between Pho and dIno80 for binding at the PRE.

## Introduction

The Homeotic (Hox) genes control body patterning in animals and are highly conserved across phyla [1,2]. The level and domain of expression of Hox genes, control segmental identity in *Drosophila* [3]. The transcription factors such as Krüppel [4] and the segment polarity genes in *Drosophila* regulate Hox gene expression during development. The maintenance of the expression is through the interaction between specific *cis*-elements, the PREs/TREs (Polycomb/Trithorax Responsive Elements) and the Polycomb (PcG) and the trithorax (Trx) complexes that constitute the Cellular Memory Module [5,6]. The proteins of the PcG complex (Polycomb Repressive Complexes PRC1 and PRC2) are highly conserved among the organisms [7]. The PcG and TrxG complexes are epigenetic modifiers that bring about heritable changes in gene expression from one generation of cells/organisms, to the next [5,8-11].

The PREs were first identified in the Bithorax complex of *D. Melanogaster [12-15].* PREs and TREs are often overlapping and contain multiple motifs for sequence-specific DNA binding proteins of the PcG/ TrxG complex [16]. In *Drosophila*, PREs are known to exert their effect on target genes that are located at long distances, ranging from a few bases to several kilo-bases [17].

The PRE/TREs contain the binding sites for Pleiohomeotic (Pho) and other recruiters of Polycomb/Trithorax complexes such as DspI, Grh, SPI/KLF, Zeste, GAGA factor, Pipsqueak and Grainyhead [11,18]. It is known that there are other factors, such as modified histones, long non-coding RNAs and short RNAs that mediate the recruitment of Polycomb/Trithorax complex [19,20]. The PRC2 member, EZH2 catalyzes the methylation of histone H3 at K27 and this modification, in turn, is recognized by PRC1 which brings about ubiquitylation of H2A at K119 leading to repression [21-23]. There are other mechanisms known for the recruitment of PcG/TrxG complex at the PREs [24-26]. As such, there are no reports of chromatin remodeling complexes interacting with PREs/TREs; however, there are a few examples where the chromatin remodeling proteins such as the Brahma (Brm) genetically interact with TrxG members. The chromatin remodeler complex further mediates transcriptional activation through their interactions with trithorax group protein Zeste [27].

dIno80 is a member of the ETP group [28,29]. The ETP members are also known to interact with PREs, for interact Corto interacts with *iab-7 PRE* [30]. dIno80/hINO80 interact with Pho/YY1 in *Drosophila* and human, respectively [31,32]. However, the interaction of this complex with PRE is not demonstrated. In *Drosophila*, the Pho-independent function of dIno80 during development and the localization of dIno80 on the polytene chromosomes in the absence of Pho is known [29]. In the present study, we provide evidence of genetic interaction and also direct interaction of dIno80 with *iab-7* and *bxd PRE*. We demonstrate that the enrichment profile of Pho and dIno80 are related to the relative concentration of each other in the cells.

## Results

### *dIno80* regulates Hox genes

In order to dissect the role of *dIno80* in development, we examined the interaction of heterozygous mutants of *dino80* with different genes. In a previous study from our lab, we demonstrated that the null mutants of *dIno80* led to alteration of Hox gene expression at the late embryonic stages [28]. The study was taken forward to examine the consequence of the genetic interaction of *dIno80* with Hox genes (*Abd-B, abd-A* and *Ubx* of the bithorax complex) in adult flies. The *dIno80* heterozygous knock-out (*dIno80*^*Δ4*^*/Sb*^*1*^) flies were crossed with Hox mutants (Fig 1 and Table1). The abnormalities occurring at high penetrance, are the fusion defects in the abdominal segments (segments 3-4 and 4-5) and altered wing margins (Table 1). Genetic interaction between *dIno80* mutant and mutants of the bithorax complex members produced progeny with abdominal fusion of segment A3 and A4 in females (Fig 1A’), an extension of the dark pigmentation in males from A5 to A4 segment (Fig1B’) and shrunken wings and brown patches on the wings (Fig 1D, E & F). Thus, suggesting that dIno80 functions as a regulator during development.

**Table 1.** Analysis of genetic interaction of dIno80 with homeotic genes. Trans-heterozygotes various homeotic gene mutations were generated in the background of dIno80 heterozygous mutation as listed. The penetrance was varied as shown.

| Genotype | Flies with altered phenotype (%) |
| --- | --- |
| *Abd-B(M1)/dIno80<sup>Δ4</sup>* | 4 |
| *Abd-B F1/dIno80<sup>Δ4</sup>* | 59 |
| *abd-A<sup>P10</sup>/dIno80<sup>Δ4</sup>* | 43 |
| *Ubx<sup>abx-1</sup>/dIno80<sup>Δ4</sup>* | 26 |
| *Ubx<sup>bxd-55i</sup>/dIno80<sup>Δ4</sup>* | 91 |
| *Antp<sup>21</sup>/dIno80<sup>Δ4</sup>* | 32 |

**Figure1.**
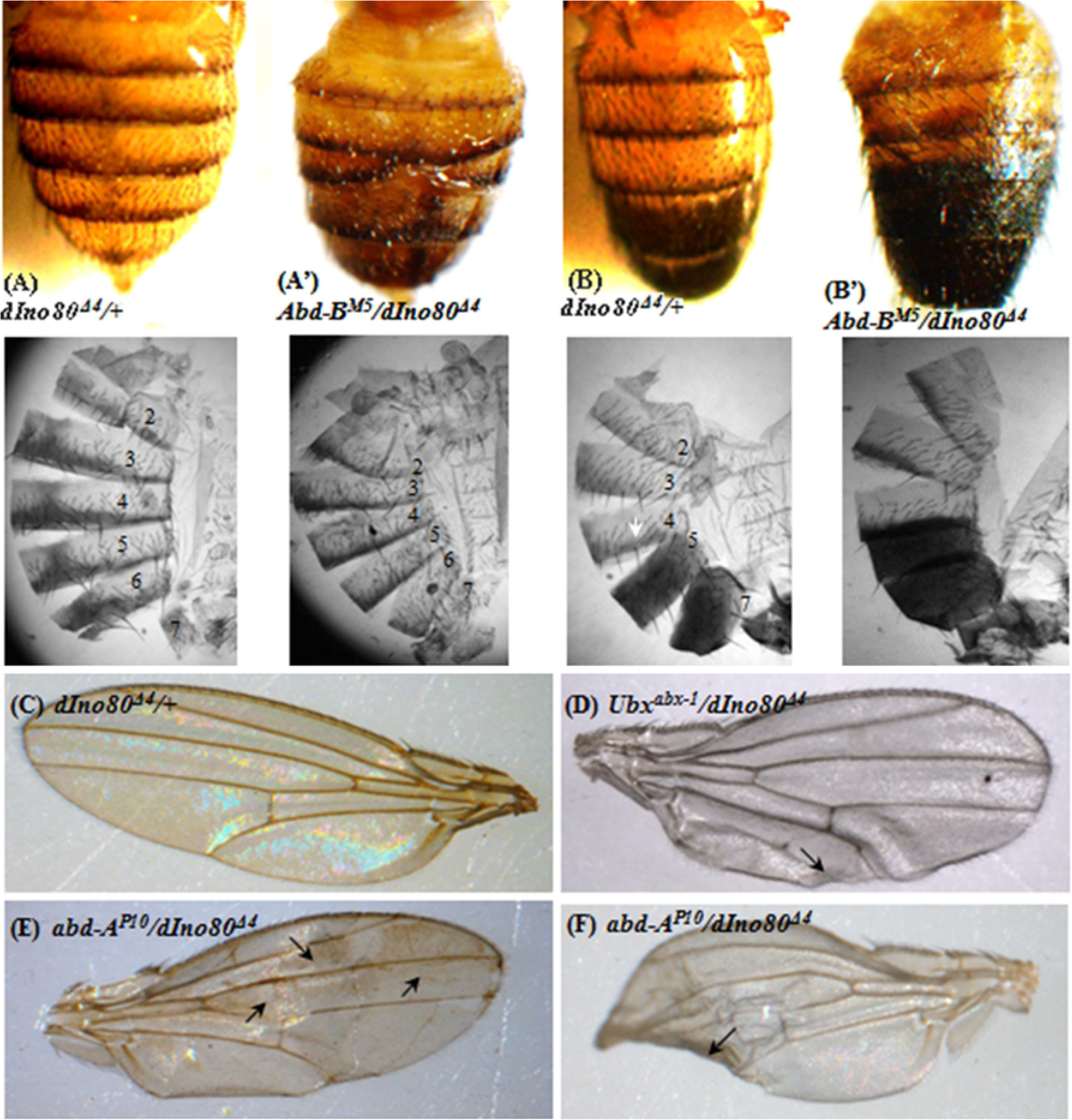
Genetic interaction of *dIno80* with Hox genes with *Abd-B, Ubx* and *abd-A*. Abdominal fusion, segmentation defects, alteration in wings are observed. The genetic background is indicated in each image. The abdominal pattern is shown in black and white images corresponding to A, A’, B and B’.

Since the regulation of Hox genes is through the PcG and TrxG complexes that interact with PRE/TRE sequences, we examined the interaction of dIno80 with PRE sequence.

### *dIno80* interacts with Polycomb Responsive elements (PRE)

We investigated the genetic interaction of *dIno80* with two known PREs of *D. melanogaster*; *iab7-PRE* that regulates the expression of *abd-A* and *Abd-B* genes and *bxd-PRE*, which governs the Ubx expression [33]. We tested different *iab-7 PRE* transgenic lines, including *wt15* and *wt44*, and observed significant changes in the expression of the *mini-white* gene in the background of *dIno80* mutation (Fig 2 and S1 Fig). Increase in the eye color in the mutant context indicates that dIno80 is required for the repressive function of the PRE.

**Figure 2.**
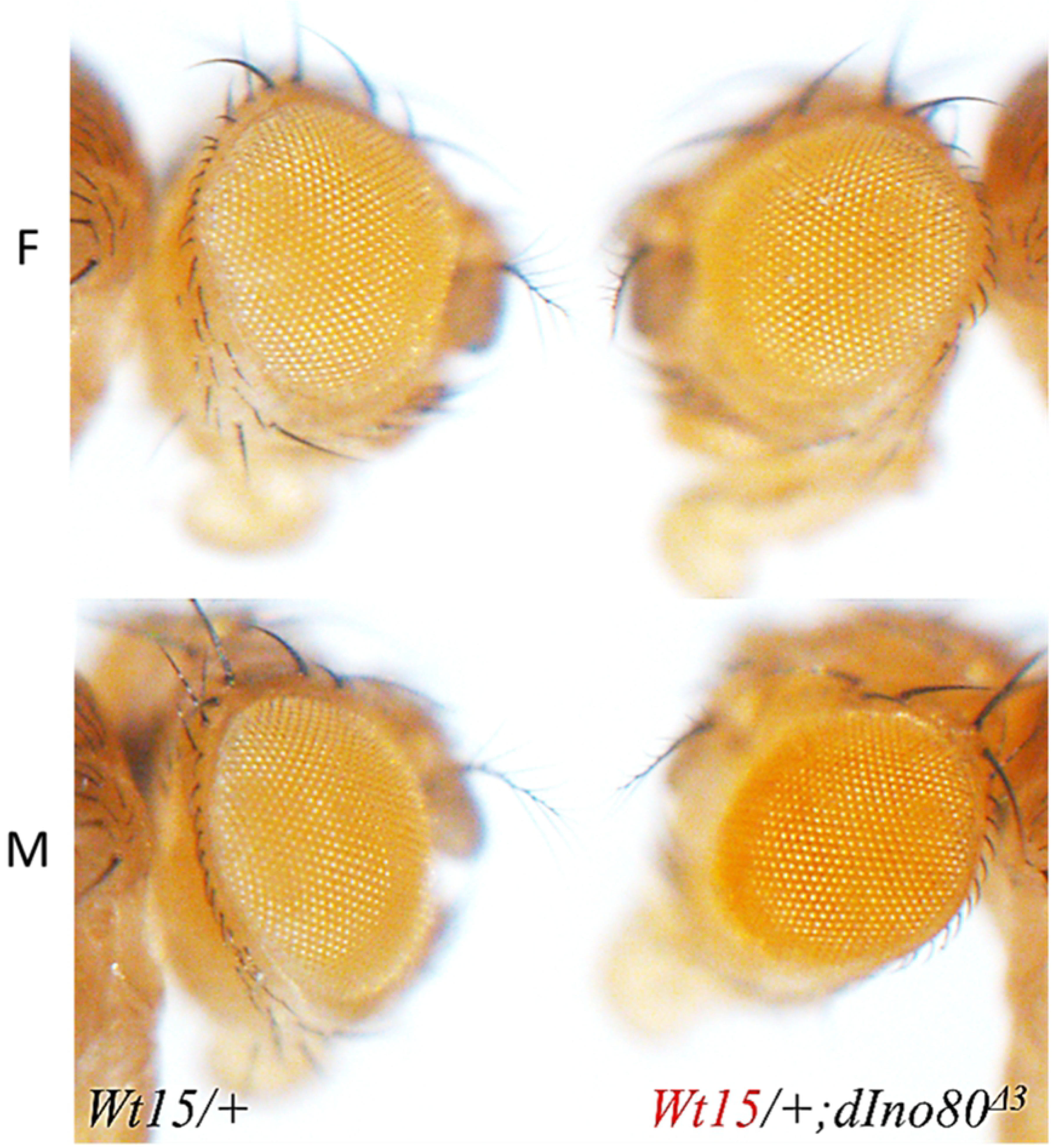
Genetic interaction of *Ino80* with *iab7-PRE (wt15)*. Mini-white gene expression is enhanced in dIno80^Δ4^ background. F-Female, M-Male. The genetic background is shown on the respective frames.

### dIno80 cooperates with PcG/TrxG complex for interaction with PRE

We studied the effect of PcG, TrxG, and ETPG mutations on the interaction of *dIno80* with the PREs. We generated double heterozygotes of *iab7-PRE* transgenic flies in the background of *dIno80* mutation along with mutants of different members of the maintenance group proteins including PcG, TrxG and the ETP group (Table S1).

We estimated the pigment content in each line following extraction from an average of 10 flies of each sex in three independent biological replicates (Fig 3 and S2 Fig). A significant difference in pigment level was observed between *iab-7-PRE* line in the background of *dIno80* mutant (*iab-7/+* ***vs*** *iab-7/dIno80*) and also between double heterozygotes of *iab-7/dIno80* TrxG, PcG and ETPG genes [(*iab-7/Trl*^*R85*^ ***vs*** *iab-7/Trl*^*R85*^; *+/dIno80*), (*iab-7/Pho*^*b*^***vs*** *iab-7/Pho*^*b*^; *+/dIno80*), (*iab-7/Asx*^*XF53*^ ***vs*** *iab-7/Asx*^*XF53*^; *+/dIno80*)]. We carried out similar assays in the background of several mutations (S2 Fig and Table S2). We have compared all the genotypes with *iab7-PRE* (*wt15*)*/+* for statistical significance (Table S2). These results suggest that dIno80 regulates homeotic gene expression through its interaction with PRE and in combination with the PcG, TrxG, and ETPG proteins. However, we cannot formally rule out the independent function of dIno80 in this regulation.

**Figure 3.**
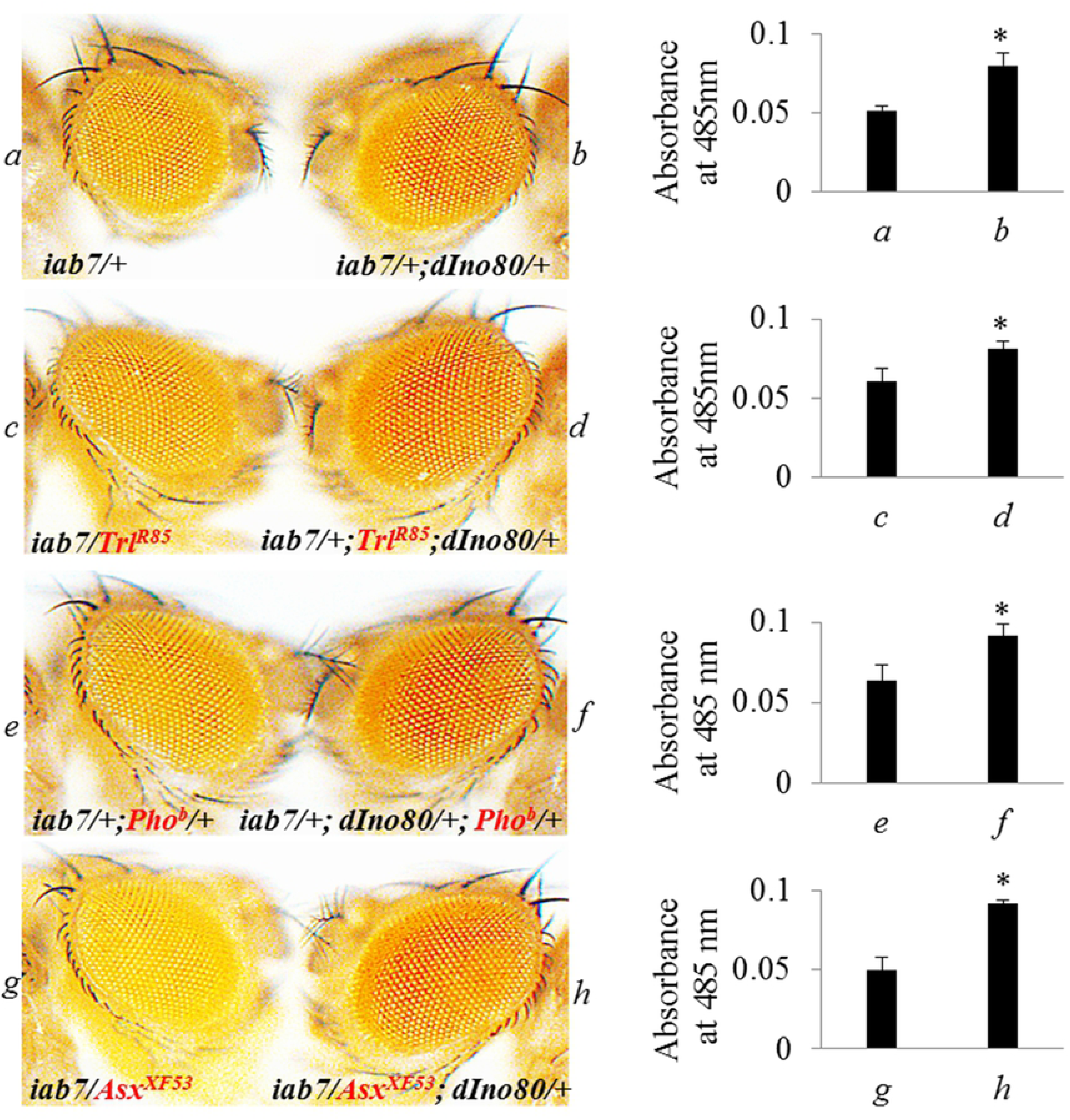
dIno80 cooperates with PcG/TrxG complex for interaction with PRE. The mini-white expression is compared between iab7-PRE (wt15) and various combinations of dIno80^Δ4^ with PcG, TrxG, and ETPG mutations. The bar graphs show the level of eye pigmentation. The mean +/- s.e.m., from three independent experiments. *p<0.05 and **p<0.001.

### Role of dIno80 in PRE mediated-regulation during development

PcG/ TrxG complexes also mediate the regulation of homeotic genes during larval stages through their interaction with PRE/TRE [34]. We examined the effect of dIno80 knockdown and deletion mutation in imaginal discs of third instar larvae of *D. melanogaster*. Since the *wt15* line with *iab-7 PRE* has only mini-white as the reporter gene, we used the *bxd-PRE* transgenic with *Lac-Z* reporter gene driven by ubiquitin promoter (Fig 4A). We compared heterozygous deletion mutant lines of *dIno80* and *Pho* (*bxd-PRE/dIno80*^*Δ4*^ *and bxdPRE/+;Pho*^*b*^*/+*) and *dIno80*RNAi lines (*bxdPRE/+;UASIno80RNAi/ ActGAL4 and bxdPRE/+;UAS PhoRNAi/ActGAL4*) for the expression of β-galactosidase in the imaginal discs.

**Figure 4.**
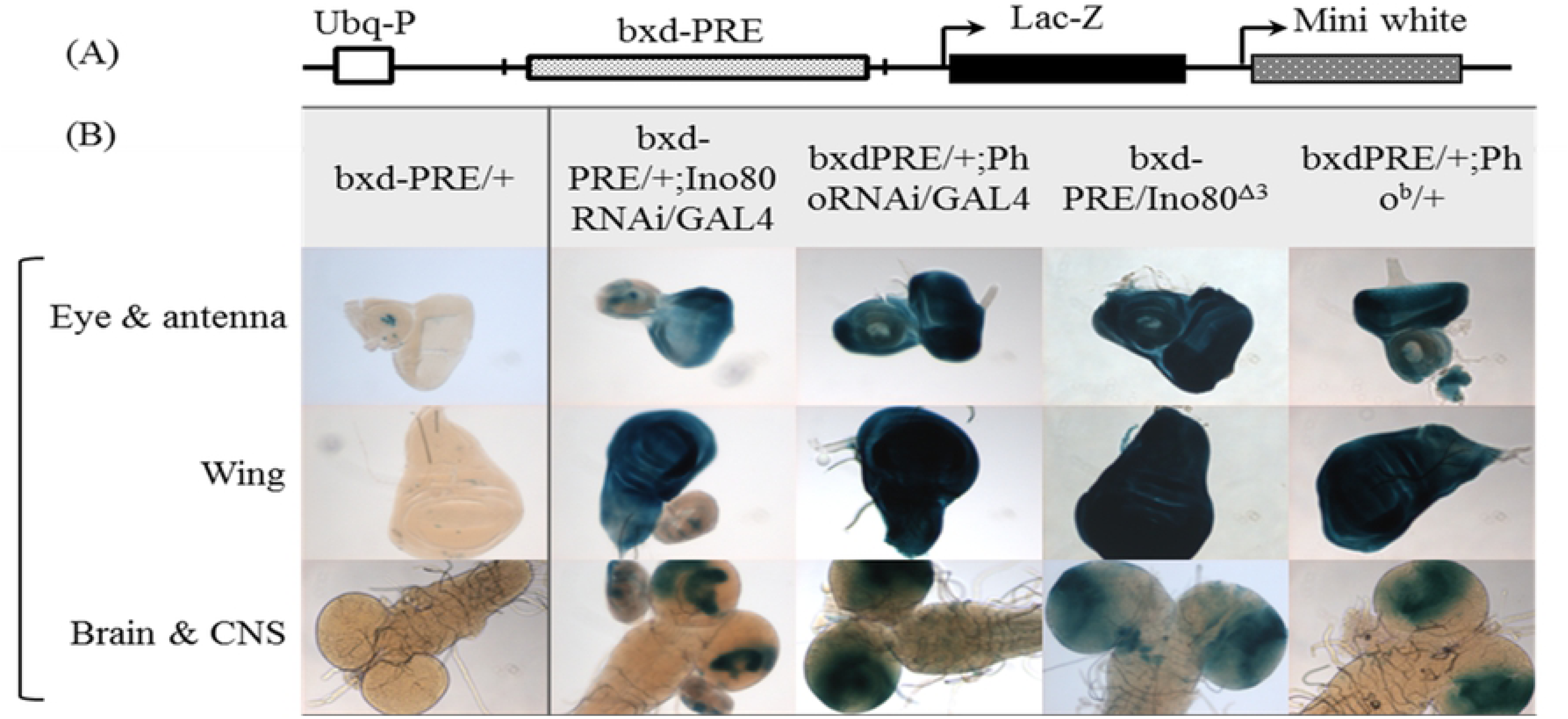
Effect of Ino80 on bxd-PRE in the imaginal disc. A-Line diagramis indicating the transgene construct. B-Lac-z expression in different imaginal discs analyzed in *Ino80* and *Pho* mutation/knock-down, as indicated. Ino80^Δ4^ is a deletion line, *Pho* and *Ino80* RNAi were driven by UAS-ActGAL4. The loss of repression is observed in both cases.

In transgenic lines with *bxd-PRE/+* in the background of *dIno80*^*Δ4*^, the repression of the β-galactosidase gene is reversed (Fig 4B). We observed a similar effect, when *dIno80* was knocked-down (*bxd-PRE: UASIno80RNAi/ActGal4)* or when *Pho* is mutated (*Pho*^*b*^) or knocked-down (*UASIno80RNAi/ActGal4*; Fig 4B). The similarity in the outcome of dIno80 and Pho depletion in conjunction with the genetic interaction of dIno80 with members of PcG, TrxG, and ETPG in iab7-PRE transgenic line strongly suggest the interaction of dIno80 with PREs through PcG complex.

Thus, we demonstrate that dIno80 regulates reporter genes (*mini-white* and *Lac-Z*) via its interaction with the *iab-7* and *bxd-PRE*, however, it needs to be elucidated whether dIno80 protein directly interacts with the PRE sequences.

### dIno80 protein is enriched at the Polycomb Response Elements (PREs)

The localization of dIno80 on chromatin has been demonstrated earlier by immunostaining of polytene chromosomes [28,29]. Since anti-dIno80 antibodies are not commercially available, we carried out the Chromatin immunoprecipitation experiments in S2 cells transfected with full-length cDNA of dIno80 with FLAG tag. The expression of FLAG-dIno80 was confirmed using anti-FLAG antibody (S3 Fig). dIno80 is known to form a complex with Pho along with other proteins [31,32]. Therefore, we investigated the localization of both Pho and Ino80 at *iab7-PRE* and *bxd PRE*.

The enrichment of dIno80 at *Bxd-PRE* (Fig 5) and *iab-7PRE* (Fig 6) was analyzed in the transfected cells using anti-FLAG antibody within the PRE and also the flanking regions. The enrichment varied between the regions. The regions flanking *Bxd-PRE* showed higher enrichment of dIno80 than the PRE itself (Fig 5). On knock-down of Pho, the enrichment of dIno80 further increased in the flanking regions while it remained unchanged within the PRE. While the knock-down of dIno80 decreased enrichment in all cases. This indicates that the binding of dIno80 is independent of Pho. The reason for differential enrichment in these regions remains to be investigated. Differential enrichment of dIno80 was observed in case of *iab-7PRE* and its flanking regions, which was sensitive to the presence of dIno80 and was enhanced in the absence of Pho (Fig 6). The reduced FLAG-Ino80 interaction on knockdown of *Ino80*, confirms the specificity of anti-FLAG antibody. The knock-down is around 80% for dIno80 and 50% for Pho (S4 Fig).

**Figure 5.**
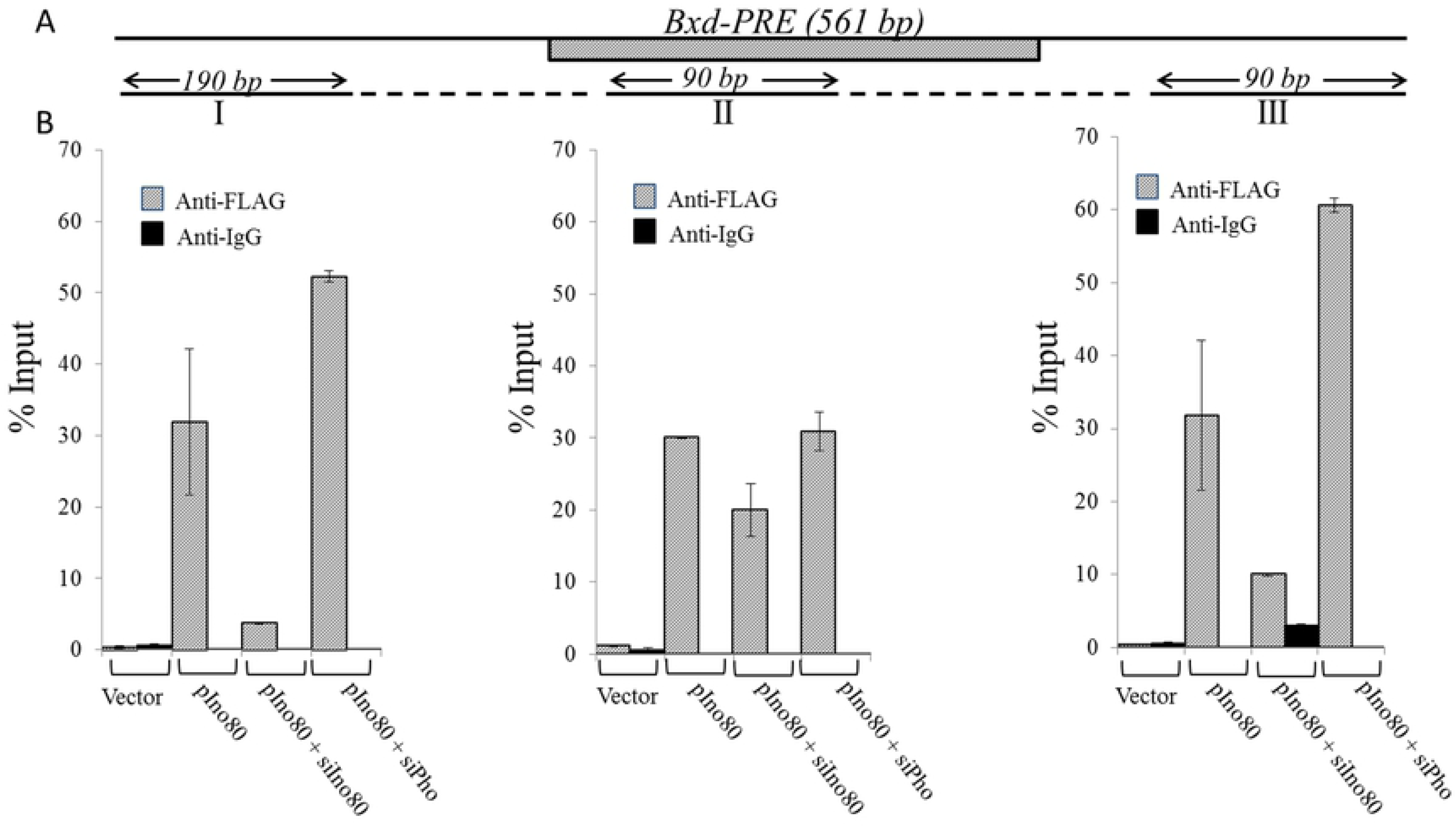
Enrichment of Ino80 on bxd-PRE. A. I-III are the regions analyzed for Ino80 binding, the amplicon size is indicated, and they span a total length of 720bp. Vector – empty pBUF. B-ChIP-qPCR assay shows the enrichment of dIno80. Anti-FLAG antibodies were used in ChIP. *p<0.05 and **p<0.001.

**Figure 6.**
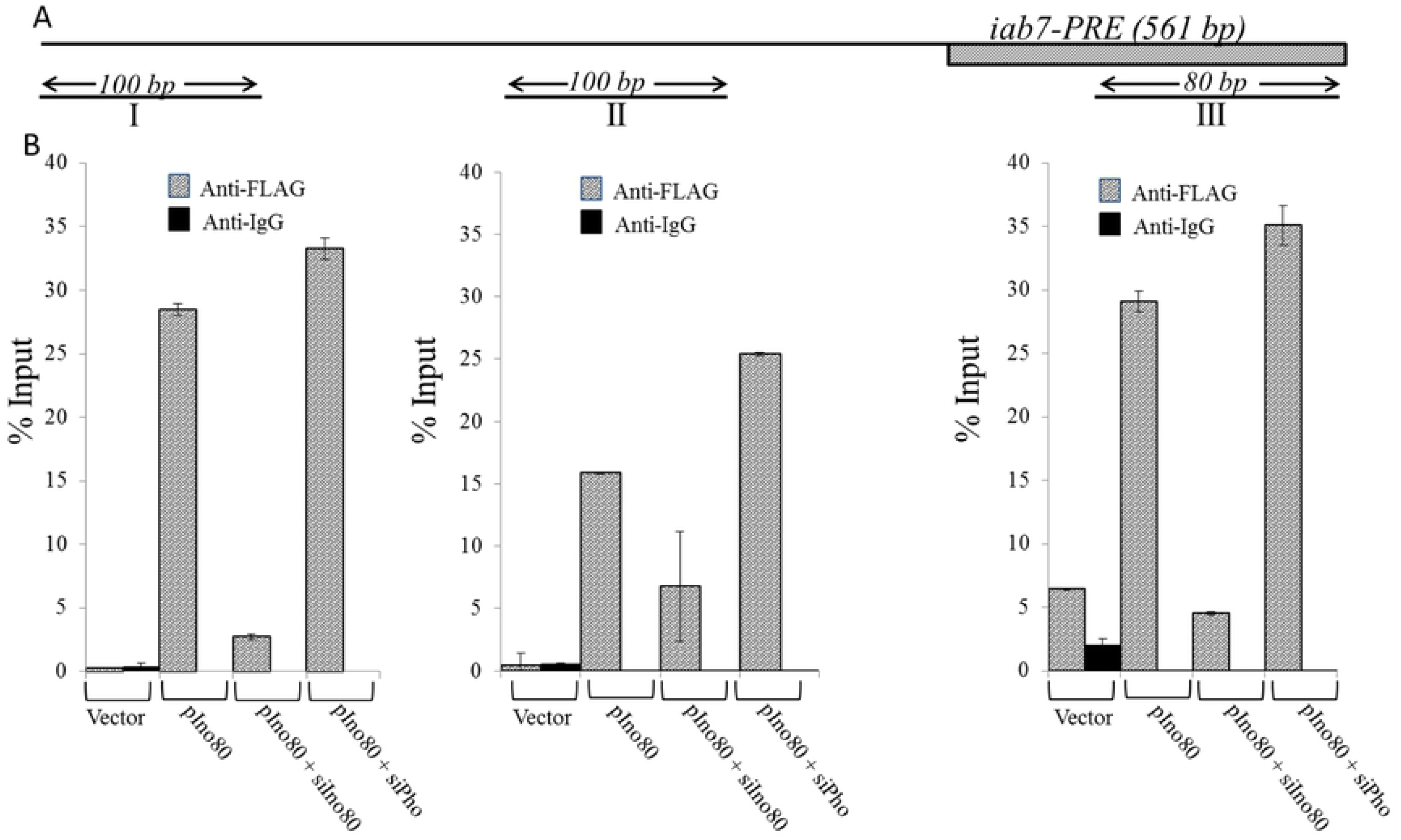
Enrichment of dIno80 on iab7-PRE. A-Indicates the regions analyzed (I-III), relative to the PRE sequence along with the amplicon size over a total length of 430bp. B-ChIP-qPCR assay shows the enrichment of dIno80. Vector-empty pBUF, and the constructs and oligos used in transfection are shown on the X-axis. Anti-FLAG antibodies were used in ChIP. *p<0.05 and **p<0.001.

The enrichment of Pho in both *Bxd-PRE* and *iab-7PRE* was assayed using Anti-Pho antibody (Figs 7A and B). On over-expression of dIno80, Pho enrichment decreases, however on knock-down of the over-expressed dIno80, the enrichment of Pho within Bxd-PRE and region downstream is observed but in the upstream flanking region it remains poor. This is consistent with the significantly high localization of dIno80 at this region when Pho is knocked down (Fig 5). In the *iab-7 PRE*, the overexpression of dIno80 decreases the enrichment of Pho in all the regions, which is partially reversed on dIno80 knock-down. These results show the Pho independent binding of dIno80, but the relative enrichment of the two proteins at the regions is not uniform across the regions examined. This may have its bearing on the existence of variable affinity of the sites known in other cases [35].

**Figure 7.**
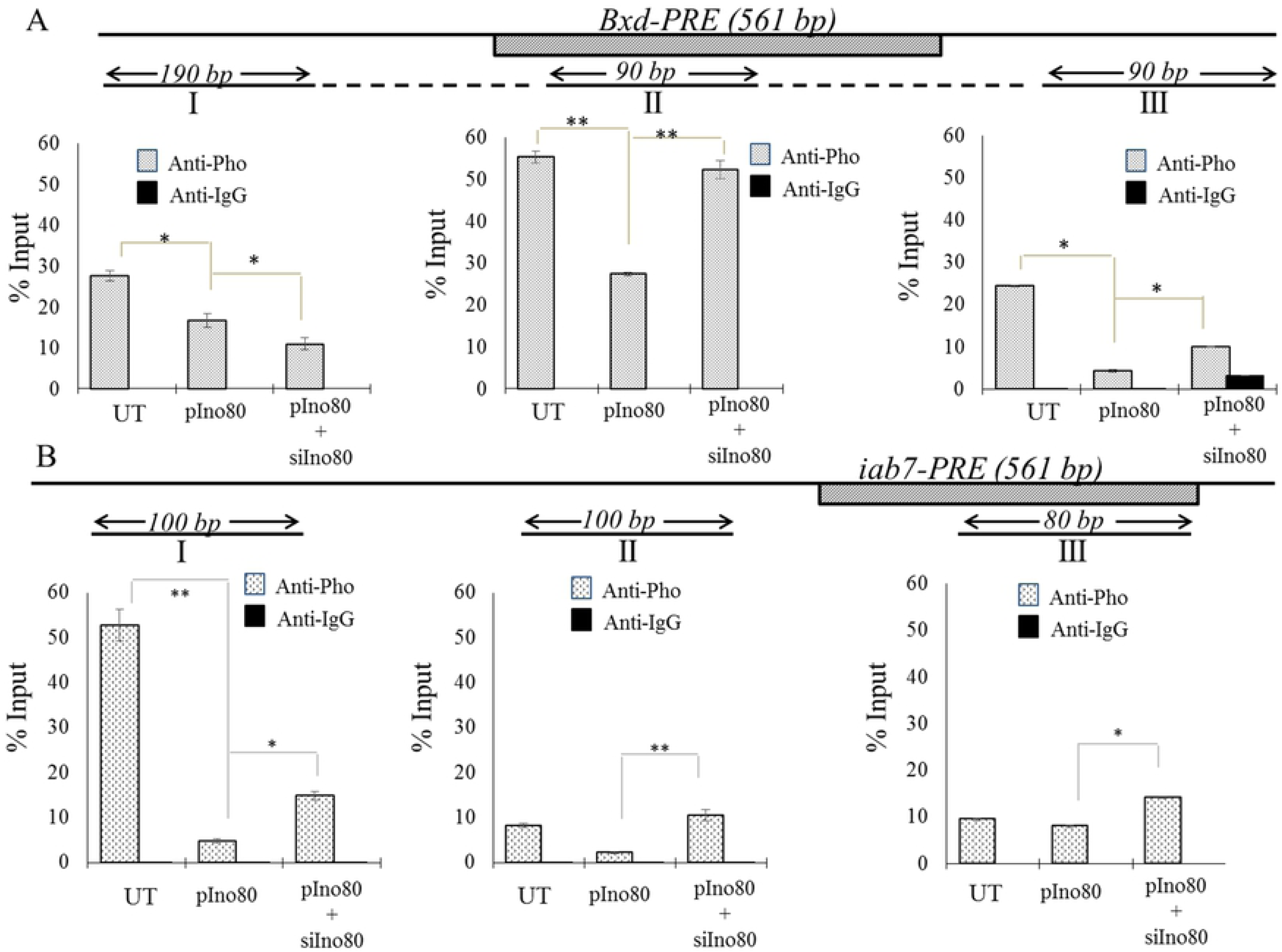
Enrichment of Pho on the bxd-PRE and iab7-PRE. A. Line diagram of *Bxd-PRE* of 561bp, I-III are the regions analyzed for Pho binding, the amplicon size is indicated, B-Enrichment of Pho shown by ChIP-qPCR. C-the regions analyzed (I-III) around the *iab7-PRE* sequence, the amplicon size over a total length of 430bp is indicated, D-ChIP-qPCR data for iab-7-PRE. UT – Untransfected and the constructs and oligos used in transfection are shown on the X-axis. The results represent mean +/- s.e.m., from two independent experiments. Anti-Pho and Anti-IgG antibodies were used in ChIP. *p<0.05 and **p<0.001.

### Functional complementation of Pho by dIno80

The lethality due to the lack of Pho and dIno80 proteins manifest at different stages of development in the fly [3], suggesting that they have developmental functions independent of each other. As both dIno80 and Pho are involved in transcriptional regulation and both are temporally differentially expressed, we examined whether dIno80 can rescue the pharate/pupal lethality caused by *pho* knockdown. The dIno80 was over-expressed in Pho knockdown background using UAS-RNAi approach with nos>Gal4 as the driver. The scheme of the cross set up is given in Supplemental Figures S5 and S6. The over-expression of dIno80 in *pho* knockdown background led to 10% rescue of pharate lethality in F_2_ and 46% rescue in the F_3_ generation (Table 2, S6 Fig). Therefore, over-expression of dIno80 early in development can partially rescue the lethality in Pho null mutants. The transcriptional regulation by dIno80 in the absence of Pho and rescue of Pho lethality may rely on the DNA binding potential of the dIno80 protein. This is supported by the localization of dIno80 on polytene chromosomes in Pho-RNAi line [3]. The over-expression of dIno80 was estimated to be 8.5-fold as compared to *Canton-S* embryos in UAS-GAL4> dIno80 line, and knock-down of Pho was 0.76-fold as compared to wild type in the UAS-Gal4 RNAi line (S7 Fig). Interestingly, the knockdown of Pho leads to over-expression of dIno80 by 2.5 fold as compared to *Canton-S*. Also, the over-expression of dIno80 led to downregulation of Pho, possibly indicating the competitive nature of binding interactions involved in transcriptional regulation (S7 Fig).

**Table 2.**
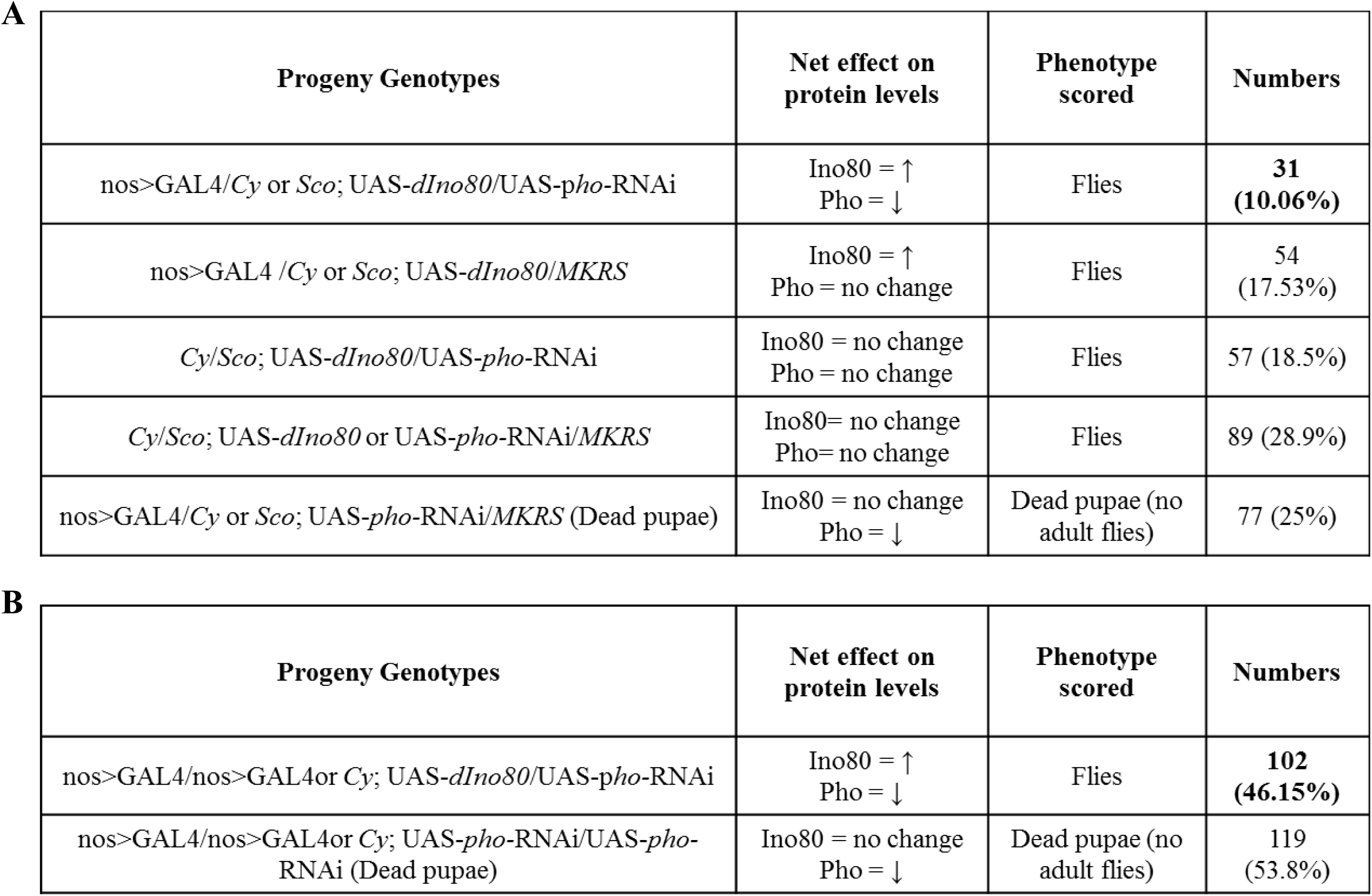
Over-expression of dIno80 rescues pupal lethality in Pho knock-down lines in F_2_generation (A) and F_3_ generation (B). The dIno80 over-expression and pho knock down was driven by nos>GAL4. The number of flies with different genotypes are shown in terms of absolute number and also as percentage.

**A**
| Progeny Genotypes | Net effect on protein levels | Phenotype scored | Numbers |
| --- | --- | --- | --- |
| *nos>GAL4/Cy or Sco; UAS-dIno80/UAS-pho-RNAi* | Ino80 = ↑<br>Pho = ↓ | Flies | **31**<br>**(10.06%)** |
| *nos>GAL4 /Cy or Sco; UAS-dIno80/MKRS* | Ino80 = ↑<br>Pho = no change | Flies | 54<br>(17.53%) |
| *Cy/Sco; UAS-dIno80/UAS-pho-RNAi* | Ino80 = no change<br>Pho = no change | Flies | 57 (18.5%) |
| *Cy/Sco; UAS-dIno80 or UAS-pho-RNAi/MKRS* | Ino80= no change<br>Pho= no change | Flies | 89 (28.9%) |
| *nos>GAL4/Cy or Sco; UAS-pho-RNAi/MKRS* (Dead pupae) | Ino80 = no change<br>Pho = ↓ | Dead pupae (no adult flies) | 77 (25%) |

**Table 2.** Over-expression of dIno80 rescues pupal lethality in Pho knock-down lines in F<sub>2</sub> generation (A) and F<sub>3</sub> generation (B).
| Progeny Genotypes | Net effect on protein levels | Phenotype scored | Numbers |
| --- | --- | --- | --- |
| *nos>GAL4/nos>GAL4or Cy; UAS-dIno80/UAS-pho-RNAi* | Ino80 = ↑<br>Pho = ↓ | Flies | **102**<br>**(46.15%)** |
| *nos>GAL4/nos>GAL4or Cy; UAS-pho-RNAi/UAS-pho-RNAi* (Dead pupae) | Ino80 = no change<br>Pho = ↓ | Dead pupae (no adult flies) | 119<br>(53.8%) |

### dIno80 affects PSS (Pairing Sensitive Silencing)

In *Drosophila*, many PREs including the *iab-7 PRE*, show Paring Sensitive Silencing (PSS) which is attributed to the somatic pairing of chromosomes and the involvement of different proteins [36, 37]. The PSS is phenotypically assayed by mini-white reporter gene expression in the eye. We tested the deletion mutant of *dIno80* for disruption of PSS in *iab-7* PRE homozygotes with Pc and Trl as positive controls. The enhancement in eye pigmentation is observed in *iab-7/iab-7; dIno80/+* flies suggesting disruption of PSS (Fig 8).

**Figure 8.**
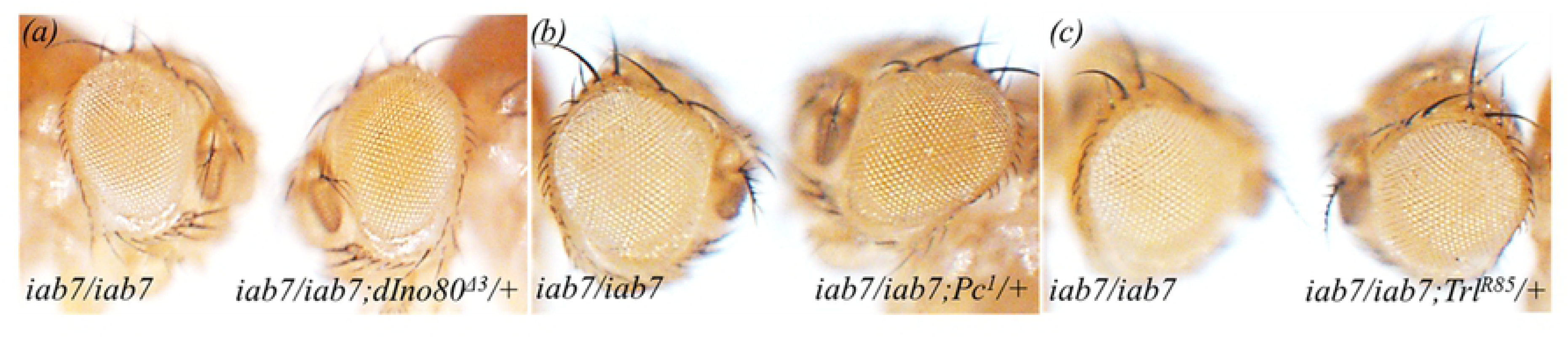
dIno80 mutation disrupts Pairing Sensitive Silencing at iab7-PRE (wt15). Image shows disruption of PSS at iab7-PRE by dIno80 (a). The loss of PSS at iab7-PRE in Pc1 (b) and TrlR85 (c) background serves as the positive control.

### INO80 interacts with *hPRE-PIK3C2B*

*hPRE-PIK3C2B* is a well-characterized PRE/TRE element that is known to interact with both the activating (TrxG) and repressive (PcG) complex members [38,39]. *hPRE-PIK3C2B* is present in the 1^st^ intron of *PIK3C2B* gene. We performed genetic interaction studies using the *hPRE-PIK3C2B* transgenic fly line *(PI-17)* in the background of *dIno80*^Δ4^ mutation and observed a decrease in eye pigmentation in comparison to *PI-17/+* (Fig. 9A).

**Figure 9.**
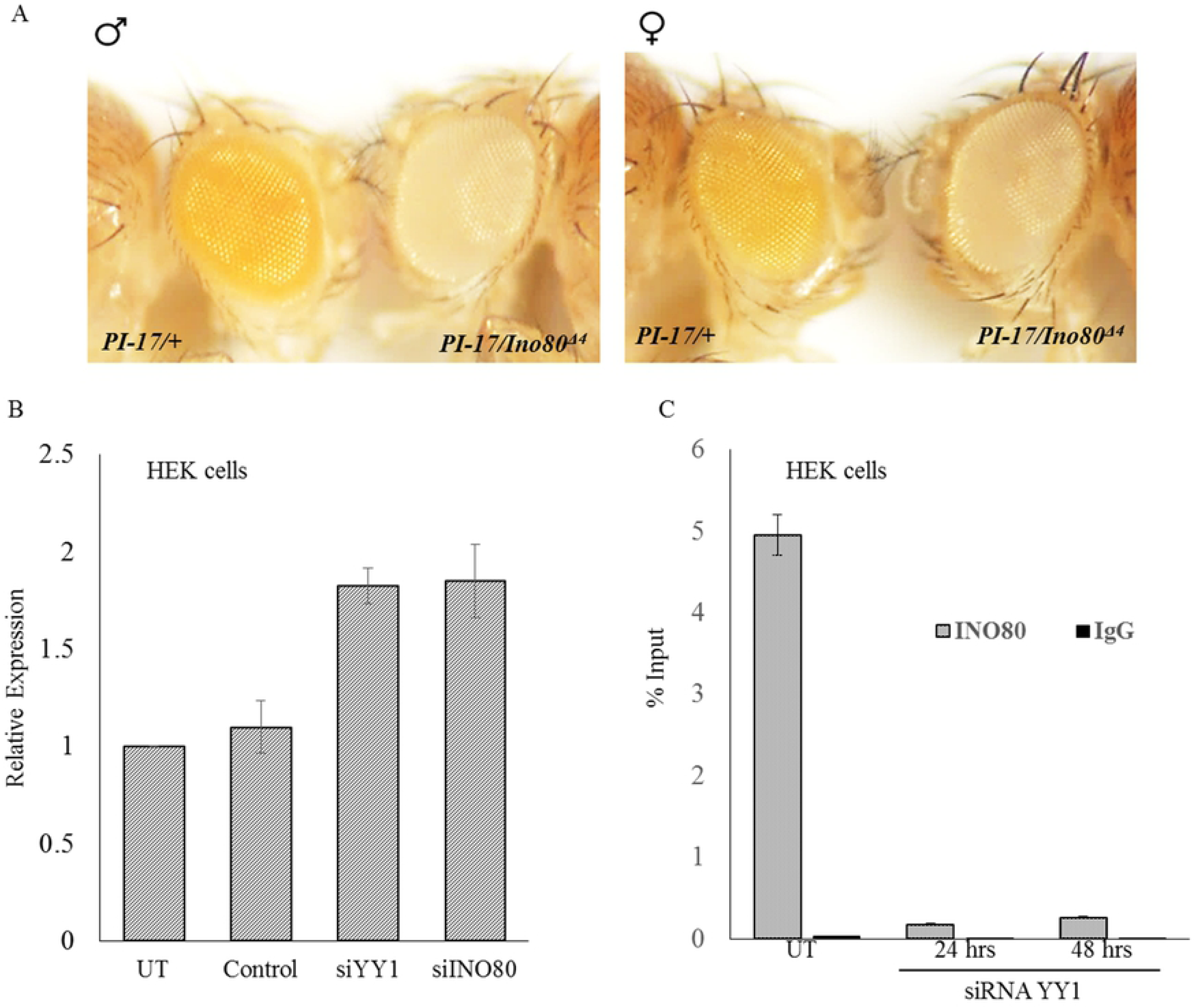
dIno80 and hINO80 interact with *human PRE-PIK3C2B.* We tested the interaction of *dIno80* and *hINO80* with human PRE in transgenic flies (A) and HEK cells (B, C) respectively. A-eye pigmentation in PI-17, hPRE-PIK3C2B transgenic line (**PI-17**) is drastically reduced in *dIno80*^*Δ3*^ mutant. In HEK cell line, knock-down of either *YY1* or h*INO80* increases the expression of *PIK3C2B* (B). INO80 interacts with *hPRE/TRE PIK3C2B*, and YY1 knockdown decreases the enrichment of INO80 at the PRE(C). Thus, YY1 mediates the recruitment of h*INO80* at hPRE-PIK3C2B in human cells.

Further, we examined the interaction of human INO80 with *hPRE-PIK3C2B* in HEK cell line (Fig 9B). The repressive effect of *YY1* (the human homolog of Pho) and *INO80* on the expression of endogenous *PIK3C2B* gene was reversed when either hINO80 or YY1 expression was knocked-down using siRNA (Fig 9B). The chromatin immunoprecipitation experiments demonstrated enrichment of INO80 at *hPRE-PIK3C2B*, which drastically decreased upon knockdown of YY1 (Fig 10C). Unlike the PREs in the fly, the human INO80 shows YY1 dependent recruitment to the *hPRE-PIK3C2B*. We have earlier demonstrated that *hPRE-PIK3C2B* has multiple repeats of YY1 binding site [38].

## Discussion

INO80 subfamily of chromatin remodelers was identified as the regulator of transcription, and the mechanism involves nucleosome re-arrangement which in turn leads to the increase in the accessibility of the chromatin [40]. The present study further adds to the already known diversity of Ino80 protein. The Ino80 protein is a versatile protein that is involved in multiple functions [29,41-44]. The interaction of dIno80 with promoters of pluripotency genes which are dependent on Oct4/WDR5 binding is also known [45]. The combination of catalytic activity of a DNA-dependent ATPase and the presence of the DNA binding domain (DBINO domain) and partnership with different proteins perhaps contributes to this functional diversity of INO80 protein in different species.

In the current study, we have shown that dIno80 directly interacts with PREs, strongly indicating that the basis of the previously demonstrated role of Ino80 in development is mediated by its interaction with chromatin [28,29]. The fusion of abdominal segments as a consequence of the interaction of dIno80 with Abd-B and also the altered wing patterning indicates the interaction of dIno80 with homeotic genes. Earlier, we reported the dual regulation of Scr, by dIno80 in different imaginal discs; being an activator of Scr in wing imaginal disc while in the leg and salivary gland disc, it acts as a repressor [29]. Thus, dIno80 interacts with several homeotic genes and based on our results, we predict that this interaction could be through the PRE/TRE sequences. The effect of dIno80 on PRE-mediated expression of the β-galactosidase reporter gene in imaginal discs through, *bxd-PRE* is very similar to that of *Pho*. This can be due to the canonical complex of Ino80 where Pho acts as the recruiter; however, the direct interaction of dIno80 with *bxd-* and *iab7* PRE even in the absence of Pho and the fact that dIno80 can rescue Pho null mutants with considerable penetrance (43%) indicates the involvement of a non-canonical complex. The genetic interaction strongly suggests that the effect of dIno80 on the transcription of genes driven by PRE sequences is through PcG, TrxG and ETPG members. Thus, as an ETP protein, dIno80 enhances the effect of Trl, Pho, and Asx on the reporter expression. It remains to be established whether Ino80 interacts with PREs directly or indirectly through other recruiters. In the light of our results, we predict the direct interaction of dIno80 with DNA.

We have shown that the DBINO domain of hIno80 as well as dIno80 bind to a consensus DNA sequence [46,47]. Therefore, we analyzed the presence of this motif and the motifs for binding of other proteins such as Pho, Dsp 1, GAF, Pipsqueak known to be involved in the recruitment of the PcG/TrxG complexes to the PREs of *D. melanogaster* (Table 3). It is observed that the putative Ino80-binding motif is present in many PREs and therefore, the direct interaction of dIno80 with PREs could be through this motif in addition to the known mechanism whereby Pho/YY1 recruits the Ino80 complex.

**Table 3.**
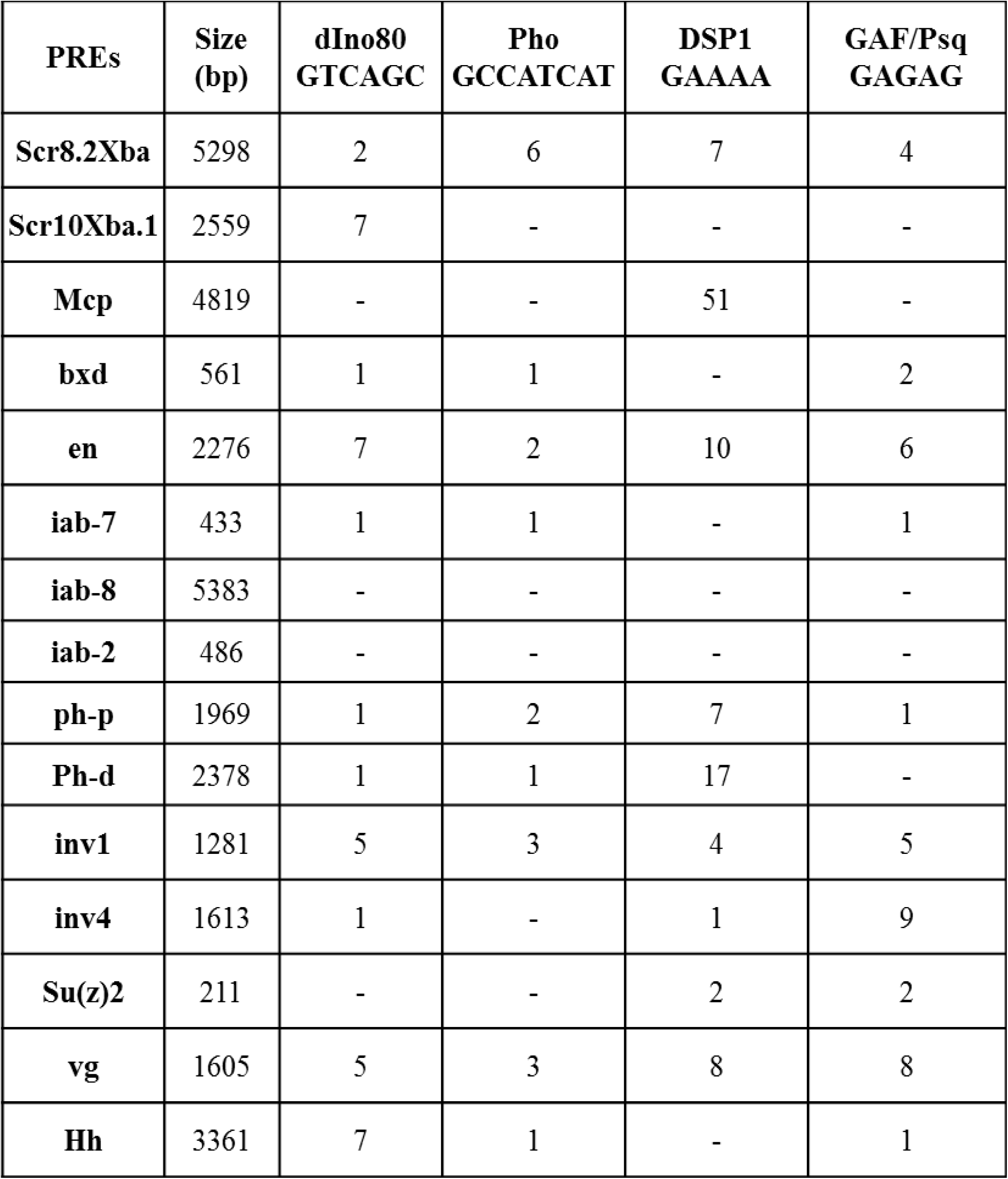
The DNA motifs within PREs of *D. melanogaster.* The size of eachPREs and the number of different motifs within the sequences including dIno80 binding site are indicated.

There are other modes of interaction of chromatin remodelers with chromatin, BPTF (bromodomain PHD finger transcription factor), a subunit of NURF complex recognizes H3K4me3, while CHD1 recognizes both H3K4me3 and H3K4me2 [48,49].

The localization of dIno80 in both the PREs and the neighboring regions is seen by the steady enrichment of Ino80 as well as Pho. The dependence of this interaction on dIno80 is indicated by the lack of enrichment on knock-down of dIno80. Interestingly, we observe that there is an increase in Pho localization on knock-down of dIno80. These results not only prove the direct interaction of Ino80 with PRE but also suggest concentration-dependent recruitment of the two proteins to the PRE sequences. In conjunction with the genetic interaction studies, it emerges that there can be the recruitment of either activating or the repressive complex through dIno80, which needs to be analyzed further. This is unlike Pho, which is known to recruit PRC2 complex at the PRE. The variable enrichment of dIno80 at the two PREs and their flanking region is similar to the variability in enrichment of PRC complex and the distribution of H3K27me3 observed in other cases where the strong and weak binding sites for PRC proteins is observed in large genomic regions of greater than 145kb [35]. It has been shown that the deletion of the major PcG regulated invected-engrailed (inv-en) gene complex does not affect regulation, which was attributed to the presence of several low-affinity PRE-like sequences in the region [35]. Even though the region we have examined is only 800-430bp, the results indicate that there is interaction beyond the limits of *Bxd* and *iab-7* PRE.

One of the important properties associated with PREs is Pairing-Sensitive Silencing (PSS). It is well known that mutations in PcG members disrupt PSS [50]. Ino80 disrupts *iab-7 PRE* associated PSS as indicated by an increase in the eye color. There are several examples of chromatin remodeling proteins participating in silencing mechanisms. The point mutations and deletions in a number of chromatin remodeling proteins and epigenetic modifiers are known to suppress position effect variegation and Telomere position effect. In concert with this mutations in SET, preSET and postSET domains of *Su (var)3-9* suppress position effect variegation (PEV) strongly, and Telomere position effect moderately [51] and mutations in *Hdac1* suppress PEV as well as TPE [51]. Chromodomain proteins such as *kis, Chd1, Mi-2*, and *CHD3* suppress PEV and TPE [51].

The developmental genes and their regulators are known to be highly conserved through evolution. dIno80 binds to hPRE-PIK3C2B in transgenic flies to bring about activation, as we observe a decrease in eye pigmentation in the background of *Ino80*^*Δ3*^ mutation. In terms of localization of hINO80 at hPRE-PIK3C2B, as measured by ChIP in HeLa cells, this interaction is completely lost when YY1 was knocked down. Thus, indicating that INO80 is recruited to *hPRE-PIK3C2B* by YY1. This is in contrast to the results we obtained where Pho and dIno80 are enriched at *iab-7 PRE* and *bxd-PRE* in the absence of the other; knockdown of *ino80* leads to increased enrichment of Pho. This strongly supports the Pho-independent role of Ino80, shown earlier in flies [29]. Previously, it has been shown that *YY1* (Human homolog of *Drosophila* Pho) knock-down led to an increase in the expression of *PIK3C2B* [38]. Here, we show that *INO80* knockdown also rescues the expression of *PIK3C2B* in HEK cells.

Our results do not formally rule out the role of ATP dependent chromatin remodeling activity of dIno80 in transcription regulation. However, the partial complementation of dIno80 of Pho function raises the possibility that Ino80 could function as a recruiter of PcG/Trx complexes. The over-expression of dIno80 can rescue Pho mutant with significant penetrance (40%), while Pho null mutation shows 100% lethality. Thus dIno80 appears to complement the function of Pho at certain developmental stages. The number of known DNA-binding recruiters coded in the fly genome is limited in the context of the diversity of developmental regulatory complexes and the sites of their recruitment. There is a report of dINO80-Pho complex, in which Pho recruits dINO80 complex to the genome [52]. Pho is essential for the completion of development in *Drosophila* and is a well-characterized polycomb group member and its knock out is pupal lethal [53]. In spite of the well-known essentiality of Pho in development, it is interesting to note that over-expression of dIno80 can rescue Pho lethality. This strongly suggests the functional complementation by dIno80 in terms of the recruitment of regulatory complexes on the genome.

Apart from its role in the development and DNA repair, Ino80 also plays a vital role in maintaining the boundary between euchromatin and heterochromatin. Ino80C blocks Dot1-mediated H3K79 methylation [54]. Thus the functional diversity of chromatin remodeling proteins can be mediated by their interaction with different protein complexes. The diversity of protein complexes formed by INO80/dIno80 protein is currently being investigated in our laboratory.

It can be hypothesized that proteins belonging to various functional classes may be brought together to achieve a unique functional outcome in a manner very similar to the concept of a “lego set.” Rvb1 and Rvb2 proteins are partners of Ino80 in the chromatin remodeling complex and are also known to interact with Tip60 complex (histone acetyltransferase), telomerase complex, snoRNP complexes and the mitotic spindle assembly [55]. Brahma, which is a part of the SWI/SNF chromatin remodeling complex is associated with MeCP2 and plays a role in transcriptional silencing [56] and DNMT3B interacts with HDAC1 and 2 along with hSNF2H (chromatin remodeling enzyme) and co-localize at heterochromatin regions [57]. Thus, proteins diverse in nature associate with each other to accomplish a novel function.

The regulatory function of Ino80 in development through interaction with PRE demonstrated here may be a tissue-specific function, and it remains to be seen whether this function is the result of a novel interaction of Ino80 either with proteins and/or RNA molecules. However, it is clear from several examples that there is a crosstalk between chromatin remodelers and well-known silencing mechanisms mediated either by Polycomb proteins or the HP1 protein.

## Materials and Methods

### *Drosophila* stocks and mutants

Wild-type (*Canton*-*S*) *Drosophila melanogaster* was used in all the experiments, UAS-Ino80-RNAi, UAS-pho-RNAi (VDRC line: UAS-Ino80-RNAi 106684, 40213 and 40214 and UAS-pho-RNAi 39529) were obtained from VDRC, Vienna Stock Centre and daGAL4, ActGAL4 flies from Bloomington Stock Center, Indiana University and were maintained at 25^0^C. dIno80^Δ4^ was generated through P-element excision in our lab and maintained against the balance rTM6Tb^1^ or Sb^1^ [28]. To investigate the interaction of *dIno80* with PRE in *Drosophila*, we set up a series of crosses with the PRE-transgenics and Polycomb, Trithorax and ETPG mutants. The fly stocks were maintained in standard corn-yeast media at 25°C. The PRE-transgenic lines, along with *dIno80* mutation, were crossed with various PcG & TrxG mutant stocks in white eye background. We utilized multiple PREs, iab7- and Bxd-PREs, mouse, and human PRE lines [38,58] for these assays. The PcG, TrxG and ETPG mutant flies were used to study the interaction of *w; dIno80*^*Δ4*^*/Sb*^*1*^ along with these PREs. PI-17 transgenic line (*hPRE-PIK3C2B*), *ino80*^*Δ3*^/Sb^1^ was used for the genetic interaction studies involving the human PRE. The different alleles used in the present study are as follows: PRC2 members (esc^2^, *Su(z)12*^*1*^, *E(z)*^*731*^, *escl*^*d01514*^, *Rpd3*^*def24*^ and*E(pc)*^*1*^), PRC1 members (*Sce*^*1*^, *Pc*^*1*^, *ph-d*^*401*^ *ph-p*^*602*^, *ph-p*^*lac*^, *ph-d*^*401*^, *Psc*^*1*^ *and Scm*^*R5-13 B*^), PhoRC members (*Pho*^*b*^*and phol*^*81A*^), ETPG members (*Asx*^*XF53*^,*Su(z)2*^*1.a1*,^ *grh*^*B37*^, *Kis*^*02*^ *and lolal*), TrxG members (*Trx*^*1*^, *brm*^*02*^, *osa*^*2*^, *ash1*^*B1*^, *ash2, Trl*^*R85*^ *and Snr1*^*01319*^) and Nurf complex (*Nurf-38*^*k16102*^). The level of mini-white expression in the progeny from different lines was assayed from 4-day-old flies of approximately the same size [59]. The genetic interaction was tested by crossing virgin females from the mutant stocks with males from the PRE-transgenic lines.

UAS-*Pho*-RNAi (VDRC line: 39529) was obtained from VDRC, Vienna Stock Centre (www.vdrc.at), UAS-*dIno80* from The Exelixis Collection at the Harvard Medical School and *nos*>GAL4 flies from Bloomington Stock Center, Indiana University and were maintained on corn agar medium at 25 °C.

### Knockdown/over-expression of genes using UAS/Gal4 system

The UAS/Gal4 system was utilized as required either for over-expression or the RNAi for the desired gene [60,61]. For *RNAi* mediated knock-down of genes, we used UAS-*Ino80-RNAi* and UAS-*Pho-RNAi* (VDRC line: UAS-*Ino80-RNAi* 106684, 40213 and 40214 and UAS-*Pho-RNAi* 39529) that were obtained from VDRC, Vienna Stock Centre and daGAL4 and ActGAL4 flies from Bloomington Stock Center, Indiana University. These stocks were maintained at 25^0^C. In all RNAi and overexpression experiments, we always used females from Gal4 lines while male from RNAi or overexpression transgenic lines to employ maternal effect. All the phenotypic quantifications in the knockdown or overexpression experiments are done using at least 40 larvae, pupae or adult flies of the selected genotype. The selection of pupae and larvae of required genotypes was based either on the loss of markers (Tubby) on balancer chromosome or Gal4 driven GFP expression.

### Immunostaining of embryos

10– 14 hours old embryos of the desired genotype were used for immunostaining as per the published protocol [62]. Anti-abd-A (Santa Cruz; sc-27063) primary antibody and Donkey polyclonal to Goat IgG - H&L (HRP) (ab6885) secondary antibody were used at a dilution of 1:100 and 1:200, respectively. In the case of HRP chromogenic reaction was performed to stain the embryo and then CNS of the embryos of the appropriate stage were dissected out, and images were captured by using Zeiss AxioScop 2.

### Cloning full-length *dIno80* in an expression vector

The full-length *dIno80* (SDO2886, from the pOT2 vector-5117 bp) was cloned in pBUF-expression vector [obtained from Jeff Sekelsky Lab, USA (DGRC, stock No. 1230)]. pBUF vector contains a 2-kb Ubiquitin promoter, which drives a moderate level of expression in all tissues. The Sequences encoding the FLAG® epitope follows the ATG corresponding to the initial codon. The full-length cDNA for dIno80 (5117 bp) was cloned in-frame using the sites in the polylinker (for more information, please see; http://sekelsky.bio.unc.edu/Research/Vectors/Vectors.html or https://dgrc.bio.indiana.edu/product/View?product=1230). The in-frame cloning was confirmed by sequencing.

### Dissecting imaginal discs β-Galactosidase assay (X-gal staining of imaginal discs)

The protocols followed for dissection and immunostaining of imaginal discs were as described by Spratford and Kumar (2014) [63].

The imaginal discs isolated from each genotype were processed within 2 hrs. The discs were washed with 1X PBS and treated with a fixative. The standard protocol was followed for staining the discs for β-galactosidase expression [53]. The stained discs are mounted in 50% glycerol and viewed under Nikon Eclipse E600 microscope.

### Eye Pigmentation Assay

Eye pigmentation was quantified as described by Wald and Allen, (1946) using male flies (∼10) 4 days after eclosion from each genotype. The flies are frozen in liquid nitrogen, decapitated manually under a dissection microscope, and heads were homogenized in 1:1 mixture of chloroform/ammonium hydroxide (0.1%). The debris was removed by centrifugation at high speed (14000rpm) for 2 mins, and the absorbance of the aqueous phase was measured in a spectrophotometer at 485 nm. The result from a minimum of three biological replicates (each of 10 flies) was analyzed. The assay for each genotype was performed in triplicate. *p*-values were calculated using the unpaired *t-*test to determine statistical significance.

### Cell culture and transient transfection

The Schneider 2 (S2) cell lines were grown at room temperature without CO^2^ as a loose, semi-adherent monolayer in tissue culture flasks and suspension in spinners and shake flasks. S2 cells were cultured in Schneider’s *Drosophila* medium (Gibco^®^, Life technologies™, Cat no. 21720-024) containing 10% FBS (Fetal Bovine Serum, qualified, US origin, Gibco^®^, Life technologies™ Cat no. 26140-079) and Penicillin-Streptomycin (10,000 U/mL-Gibco^®^, Life technologies™ Cat no. 15140-122) at a final concentration of 50 units penicillin G and 50μg streptomycin sulfate per milliliter of medium at 25°C in the incubator (Memmert™ IPP55).,

*Drosophila* S2 cells were transfected with the recombinant expression vector for transient expression studies following the protocols described in [64]. The plasmid to transfection reagent ratio used [Attractene Transfection Reagent-QIAGEN^®^ (cat no. 301005)] was in accordance with the manufacturer’s protocol. Both plasmid and Attractene reagents were mixed separately in 500μL of the medium, followed by incubation for 5 min at RT. The cells were incubated for 24h post-transfection before further analysis.

The expression of dIno80 (full-length *dIn80* in pBUF vector) was analyzed by Western blotting using anti-FLAG^®^ antibody (Sigma Aldrich^®^, cat. No. F7425). The band intensity was measured using densitometry.

### Knock-down of *Pho* and *dIno80* in S2 cells

For carrying out the knock-down of *Pho* and *dIno80* in S2 cells, we have followed the protocol described by Celotto and Graveley (2004) [65]. In brief, we used two different pools of duplex siRNA oligonucleotide (*Eurogentec*^*©*^, Belgium), each for *dIno80* and *Pho*. We performed transfection experiments using three different concentrations for each of the siRNA pool (100, 150 and 200 pmol) for 24h and 48 h post-transfection for each gene.

### Chromatin Immunoprecipitation (ChIP) assay

The ChIP assay was performed as described by Kuo and Allis (1999) [66]. Approximately 8.0 × 10^6^ Schneider 2 (S2) cells were taken and were cross-linked at a final concentration of 1% formaldehyde G and glycine was added to a final concentration 0.125mM to quench the cross-linking. The samples were sonicated in Bioruptor (Digenode; Bioruptor TM UCD-200) at high Power; Pulse: 30 sec ON/30 sec OFF; Total Pulses 90 on ice bath) to obtain fragments ranging from 100bp to 500bp. The sonicated sample was spun at 13000 rpm for 10 minutes at 4ºC and stored at –80ºC until further use. Following the pre-clearing of sonicated chromatin using protein A/G beads (Santa Cruz sc 2003-Lot # E0115) for 4 hrs at 4°C under constant rotation, the aliquots of chromatin (25 µg) were incubated with 5µg of appropriate antibody or IgG overnight at 4°C with continuous rotation. We used immunoglobulin G (IgG) in all the Immunoprecipitation experiments as a negative control. We used anti-FLAG antibody (Sigma, cat no. F7425) for immunoprecipitation of the Flag-tagged dIno80 protein bound to the chromatin. We utilized the empty vector (pBUF vector) and un-transfected cells as negative controls in ChIP experiments. The complexes were incubated with protein A/G beads and the ChIP DNA was purified using Chelex beads (cat. No. #142-1253; BIO-RAD).

Following purification of DNA, a relative abundance of each protein at target sequence and at control regions was estimated by qPCR, using FastStart Universal SYBR Green Master (Rox) from Roche Diagnostics using ABI 7300, (Life Technologies Pvt.Ltd). The oligonucleotides (primers) used in the qPCR, which mapped to the PRE regions, are given in Table S3. The siRNA-duplex sequences used for knockdown of *dIno80* and *Pho* in S2 cell are provided in Table S4.

### Quantitative PCR

qPCR experiments were carried out in an Applied Biosystems qPCR machine, ABI 7300, (Life Technologies Pvt.Ltd), using FastStart Universal SYBR Green Master (Rox) from Roche Diagnostics GmbH, with ChIP-DNA or cDNA as template. The oligonucleotides (primers) used in the qPCR, which mapped to the PRE regions, have been listed in Appendix-IV. We analyzed the relative gene expression, using the 2^-ΔΔCT^ method [67] and the interaction with PREs using percentage input method (Invitrogen and Sigma protocols).

### Pho rescue experiments

The flies of appropriate genotypes were generated, as shown in the Supplementary figures S6 and S7. To estimate *pho* knockdown and *dIno80* expression levels, the total RNA was isolated by using TRIzol (Invitrogen) from *Drosophila melanogaster* embryos of appropriate genotype. 50-100 embryos were collected on sugar agar plates (1.5% sucrose, 3.5% agar) after about 16-20 hours post placing the flies for egg-laying (stage 14-17 of embryogenesis). The RNA was isolated using the manufacture’s protocol, cDNA synthesis, and quantitative PCR was carried out as described earlier. The primers used are *ino80*Fp: 5’-GGGAACGTCTTCACCCCTAC-3’,*ino80* Rp:5’-CAGGACACTCCAGGCGATTG-3’,*rpl32*Fp:5’-AGCATACAGGCCCAAGATCGTGAA-3’,*rpl32* Rp:5’-TCTGTTGTCGATACCCTTGGGCTT-3’, *pho* Fp: 5’-TGCACACTCACGGTCCTCGAGT-3’, *pho* Rp: 5’-TCACCGGTGTGAACCAACTGATGTC-3’. This was performed in two biological replicates each in triplicate.

## Acknowledgments

The authors are grateful to Dr. Akanksha Verma for her help with the fly images. This work was supported by the Department of Biotechnology, Govt India (BT/PR4442/PID/6/626/2011) and the Council for Scientific and Industrial Research (CSIR), Government of India (EpiHeD: BSC0118/2012-17) and No.60 (0102)/12/EMR-II). We acknowledge DBT Bioinformatics facility at ACBR and the UGC SAP-II support. Financial support through Research Fellowship for MG under DBT project (BT/PR4442/PID/6/626/2011), SJ from CSIR, Government of India, JM from SERB as Research Associate fellowship (EMR/2016/000653/dt 15/03/2017) is gratefully acknowledged.

